# Steroid hormone induced expression of LIF mediate vasoactive molecules during window of implantation and decidualization in golden hamster

**DOI:** 10.64898/2026.05.25.727212

**Authors:** Randhir Kumar, Chandana Haldar, Pranab Lal Pakrasi

**Affiliations:** Department of Biosciences, School of Science, Indrashil University, Mehsana, Gujarat-382740, India; Department of Zoology, Banaras Hindu University, Varanasi-221005. India

**Keywords:** Implantation, LIF, Letrozole, COX-2, Prostaglandins

## Abstract

Embryo implantation is early and complex stage of pregnancy begins when competent blastocyst makes a physiological attachment to receptive endometrium. Expression of numerous molecules are essential for initiation of pregnancy. leukemia inhibitory factor (LIF) is essential cytokines required for priming uterus to make it receptive for implantation. In mice, the ovarian estrogen regulated expression of LIF is absolutely required for implantation. Golden hamster showed ovarian estrogen independent process of embryo implantation. Hence, the regulation of LIF in uterus of golden hamster during early pregnancy is still ambiguous. In this study, we explored the possible regulation of LIF by uterine factor and their spatio-temporal localization and expression in the uterus of golden hamster during early pregnancy and pseudopregnancy. We further demonstrated their ability to activate prostaglandin synthesizing enzymes to achieve successful pregnancy. We used immunohistochemistry, quantitative and semiquantitative PCR to achieve the objectives. We observed the expression of LIF in all the day of early pregnancy and pseudopregnancy in the uterus of hamster. Their m-RNA was found to be upregulated around the day of implantation and decidualization. LIF showed high expression in D3 pseudopregnancy. LIF was found to be regulated by estrogen in ovariectomized uterus and significantly reduced expression of LIF was observed in letrozole treated uterine horn. Downregulated expression of prostaglandin synthesizing enzymes was observed in anti-LIF antibody treated uterus. Together, these findings highlights that uterine factor regulated LIF mediate their action via activating prostaglandin synthesizing enzymes to make uterus receptive for successful early pregnancy in hamster.

**Highlight:**

- Expression of LIF in uterus during pregnancy in golden hamster is independent from the presence of blastocyst
- LIF is regulated by estrogen in ovariectomized hamster
- Expression of LIF mRNA is downregulated in letrozole treated uterine horn in day 5 of pregnancy indicating the possibility of their regulation by uterine estrogen in golden hamster
- Prostaglandin synthesizing enzyme and LIF might be associated with the activation of inflammatory signals which are essential for successful establishment of early pregnancy in golden hamster.

## 1. Introduction

Infertility is one of the devastating problems worldwide in recent decades which adversely affect the socioeconomic status of a person. Loss of early pregnancy due to failure in embryo implantation is considered as one of the major causes of infertility in females and most important limiting factor for achieving a successful outcome using the assisted reproductive techniques (ART) (Noewitz et al., 2001; Zhang et al., 2013). Statistic report suggests that one among each nine couple in Europe and USA is affected by early pregnancy disorders related to implantation failure (Krussel et al., 2003). Embryo implantation is one of the most crucial steps in the female reproduction in which the competent blastocyst get adhered to receptive endometrium during a narrow window of time to established successful pregnancy (Ma et al., 2003). This requires a cross talk between blastocyst and endometrium to achieve a synchronized growth and development of blastocyst and attaining a stage of endometrial receptivity for successful implantation. In response of implanting blastocyst, uterus undergo extensive remodeling in which stromal cells surrounding the implanting blastocyst proliferate and differentiate in to a specialized cell types commonly known as decidual cells by a process of decidualization (Cha et al., 2012). Several implantation specific molecules like vasoactive mediators, growth factors, cytokines and transcription factors are recruited by either endometrium or/and blastocyst at feto-maternal interface during window of implantation and play very critical role in determining endometrial receptivity (Dey et al., 2004; Simon et al., 2000; Singh et al., 2011).

Leukemia inhibitory factor (LIF) and vasoactive mediators which includes prostaglandin (PGs) and nitric oxide (NO) ranked superior by implantation biologist due to their crucial role to modulate the endometrial receptivity. All these molecules are supposed to involved in the process of embryo implantation in mice and rats and are strictly regulated by ovarian estrogen and progesterone. Golden hamster is a unique model for the study of embryo implantation because they do not require ovarian estrogen for the successful event of implantation. This phenomenon is very similar to human being in which luteal phase estrogen does not play any significant role in the establishment of early pregnancy (Reese et al., 2008).

Amongst these LIF has comprehensive role in the process of embryo implantation and decidualization of stromal cells. It is pleiotropic and highly glycosylated proinflammatory cytokines belonging to IL-6 family having three spliced variants acting as paracrine factors in the attachment of blastocyst in endometrium (Tawfeek et al., 2012). LIF is responsible for modulating a number of growth factors, cytokines, cell adhesion molecules and transcription factors directly or indirectly in luminal epithelia and stroma respectively through at least 25 signaling pathway to establish and maintenance of successful pregnancy (Rosario et al., 2016). Action of LIF is mediated by LIF receptor, LIFR and gp130 which on activation transformed in to high affinity functional receptor complex that induce STAT (signal transducer and activator of transcription) as a downstream signaling pathway (Sun et al., 2013). Some reports are also available which explain the crucial role of gp130 and STAT along with LIF in early pregnancy. Null mouse of STAT-binding site and gp130 is infertile indicating their vital role in the regulation of LIF and thus, in early pregnancy (Sun et al., 2013). The expression of LIF in mice is found to be upregulated during window of implantation and stromal cell differentiation that occurs immediate after implantation, reflecting their critical role in embryo implantation and decidualization (Song and Lim, 2006). In addition, the basal level of LIF and gp-130 has been observed to express in luminal epithelial of mice throughout of early pregnancy which further emphasize their role in the process of embryo implantation (Bhatt et al., 1991; Shen et al., 1992).

Since, the hormonal milieu during embryo implantation in golden hamster is different from rat and mice. Therefore, the expression and regulation of LIF in golden hamster might differ from other mammals. Thus, the present study were undertaken to decipher the spatiotemporal expression and precise regulation of LIF in the uterus of early pregnant and pseudopregnant golden hamster. Simultaneously, we have also checked the possibility of regulation of LIF by uterine factor and their synergistic role with prostaglandin to mediate the process of early pregnancy in golden hamster.

## 2. Materials and Methods

All reagents were purchased from Sigma Aldrich, St. Louis, MO, USA unless otherwise indicated.

### 2.1 Animals care and maintenance

Golden hamsters were kept in a controlled environment at 14 hrs light and 10 hrs dark cycles in an animal house facility of Department of Zoology, Banaras Hindu University, Varanasi, India. They were provided unlimited access of food and water. 10–12-week-old hamsters were considered for mating. All experimental procedure were conducted according to the institutional guideline and strictly followed the rules of experimental animals (Scientific Procedure) Act 2007, of the Committee for the purpose of Supervision and Control on experiments on Animals (CPSCEA), Government of India, on animal welfare.

### 2.2 Sample collection

Adult female hamsters with four consecutive estrous cycles were kept with proven male on the evening of proestrus in 1:1 ratio to get dated pregnancy. Confirmation of sperm in vaginal smear in the next morning was considered as day 1 (D1) of pregnancy. Whole uterus was collected from day 1to day 3 of pregnancy and embryo were flushed from the end of oviduct using normal saline. Implantation and inter-implantation site were separately excised from the uterus of day 4 to day 8 pregnancies. Tissues were immediately stored at – 80°C for real time PCR and Western blot analyses. Some parts of tissues were fixed in bouins’s fluid for immunohistochemistry. Every experiment was performed in minimum five independent biological and technical replicates.

### 2.3 Vasectomy of male hamster for making pseudopregnancy

Male hamsters were anaesthetised using anaesthetic ether and positioned on their back for the exposure of the abdomen. Later than testes were propelled into the scrotal sac by applying moderate pressure to the abdomen. A 10 mm midline skin incision was made. To make vas deferens visible, a 5 mm incision on the left of the midline was made and pushed the testis smoothly to the left. Isolated vas deferens was held with the forceps and cut by scissors. Same procedure was executed for the right vas deferens and incision was closed with suture thread. After 2 weeks, the vasectomized males can be used for mating. Normal female hamsters were caged with healthy vasectomized male for getting pseudopregnancy.

### 2.4 Intrauterine administration of anti-LIF antibody and Letrozole

On the day 3 of pregnancy, female hamsters were anaesthetised with anaesthetic ether. Ovary, oviduct and tubal end of the uterine horn were uncovered via a dorsolateral incision through the body wall. Peritoneal cavity situated was gently removed from the cavity to exposed ovary, oviduct and a small amount of the upper part of the uterus. The top of the uterus was held gently and anti-LIF antibody (2µg/10µl, Santa Crutz, SC-1767) and letrozole (20 mg/kg body weight, Abcam/ab141293) were delivered using hamilton syringe. The uterus, oviduct and ovary were placed back into the peritoneal cavity. The body wall was stitched with thread. This procedure was repeated for the other side.

### 2.5 Immunohistochemical staining

Immunohistochemical staining was performed following method of Pakrasi and Jain, 2008 with slight modification. Briefly, the whole uteri/implantation and inter-implantation sites were removed and fixed in Bouin’s solution overnight. Further, tissues were dehydrated in ascending grades of ethanol and cleared in xylene and embedded in paraffin wax. A 7 μm tissue sections were mounted on poly-L-lysine coated slides. After deparaffinization in xylene and rehydration in descending grades of ethanol, tissues were subjected for heat induced antigen retrieval for 15 min in microwave oven using 10 mm sodium citrate buffer (pH 6.0). The slides were allowed to cool at room temperature and then incubated with normal horse serum for 2 h to block any nonspecific reaction. The sections were incubated with primary antibody of LIF (Santa Crutz, SC-1767 ) for overnight at 4°C. This is followed by incubation with appropriate biotinylated secondary antibody for 1 hr at room temperature. Endogenous peroxidase activity was blocked by incubating sections in 3% hydrogen peroxide (H2O2, Sigma, USA) in methanol for 2 min at room temperature. Slides were washed thoroughly in PBS (pH=7.4) between incubation. Finally, sections were stained using ABC kit (Vector laboratory, USA) according to the manufacturer’s instruction. The signal reaction was performed by DAB (Sigma, USA) followed by counterstaining with hematoxylin for cytoplasmic proteins before mounting. Brown deposits indicate the positive signals. Images were captured by Leica DMIRB, Leica Microsystems Wetzlar, GmbH, Germany having external camera of canon (Japan).

### 2.6 Gene expression analysis

Total RNA was extracted from the uterine horn of pregnant golden hamster using Trizol-reagent (Invitrogen, USA). RNA was treated with DNase (Ambion, USA) to avoid any DNA contamination. RNA was run on 2% gel to check their integrity and quantified by nanodrop spectrophotometer (Nanodrop Technologies,). RNA samples with 260/280 ratio ranges 1.9-2.1 was taken for further experiment. The cDNA was synthesized following manufacturer’s protocol by using commercial c-DNA synthesis kit (Bio-Rad, USA).

We performed quantitative real time PCR analysis and reverse transcription PCR for all of the genes used in study along with β-Actin as control. Primers were designed using IDT-Primer Quest with default parameters. The primer sequences are listed in table 1 and table 2. Specificity of the primers was analyzed using BLAST tool of NCBI. Real time was conducted on 96 well plate and performed on iQ5 thermo cycler (Bio-Rad, Hercules, CA, USA) with iQ5 Optical System Software version 2.0 (Bio-Rad, Hercules, CA, USA). Each reaction mixture of 20 μl contained 5 μl first strand cDNA (100 ng) as template, 0.8 μl of each gene specific primer (200 nM) and 10 μl SYBR Green qPCR Mix (Bio-Rad, USA). Quantitative RT-PCR was performed with the initial incubation for 1 min at 95°C, followed by 40 cycles at 95°C for 15 s and extension at 60°C for 60 s. The dissociation curves were achieved for the assurance of absence of nonspecific amplifications. Five biological and three technical replicates were taken for each sample, and standard deviation and standard error were calculated. As an endogenous control, β-Actin was used for the normalization of Ct value obtained and the analysis of relative expression of mRNA was performed using ΔΔCt method (Livak and Schmittgen, 2001).

**Table 1.**
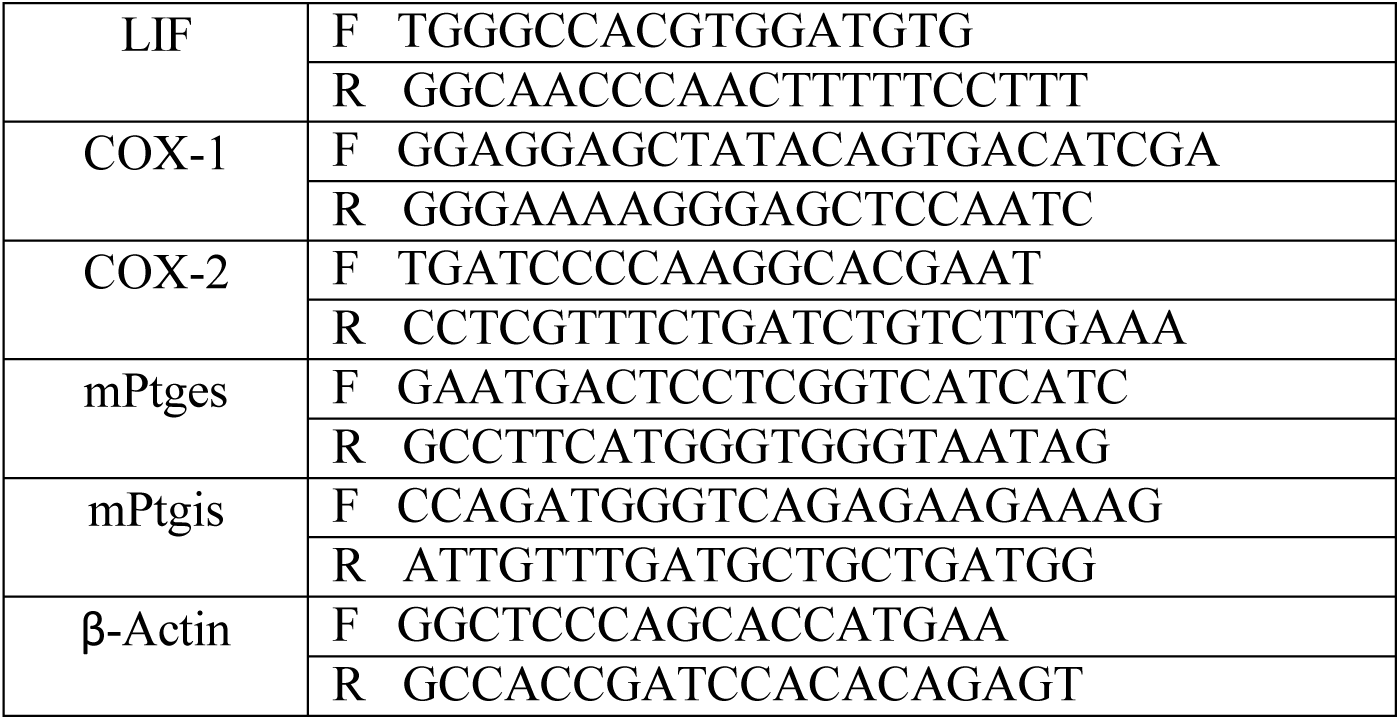
List of primers used for quantitative reverse transcriptase PCR.

**Table 2.**
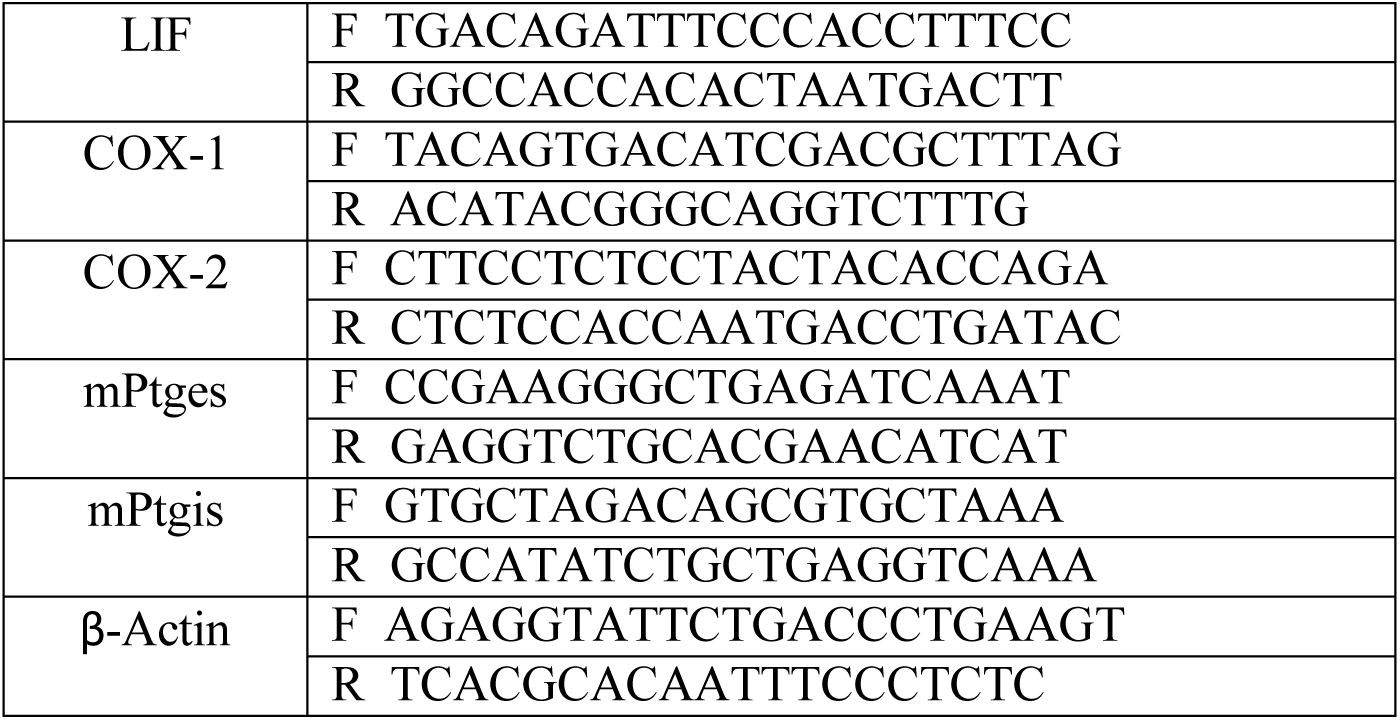
Li s t of primers used for semi-quantitative reverse transcriptase PCR.

To analyze the gene expression by reverse transcription PCR, 1 μg of total RNA was subjected to reverse transcription reaction using SuperScript II reverse transcriptase (200 units per reaction; Invitrogen, USA) with oligo(dT) primer for 50 min at 42°C. PCR amplification was performed as follows: initial denaturation at 94°C for 2 min followed by 27 cycles of incubations at 94°C for 20 s, 55°C for 40 s, and 72°C for 1.0 min, with a final extension at 72°C for 10 min with iTaq DNA polymerase (iNtRON Biotechnology, Korea) on BioRad C 1000 thermal cycler. β-Actin expression level was used as a quantitative control. Aliquots of individual PCR products were resolved in agarose gel electrophoresis and visualized with ethidium bromide under UV light. All experiments were repeated at least three times, and results from one representative experiment are shown.

### 2.6 Hormonal treatment in ovariectomized hamster

To determine the hormonal regulation of LIF, ovariectomized model were used. A total 5 hamster per hormonal treatment were ovariectomized to remove the source of internal estrogen and progesterone. All animals were allowed to recover from surgery for two weeks. Afterward, subcutenious injection of estrogen (1 µg/hamster, sigma-aldrich), progesterone (500 µg/hamster, Sigma aldrich) or a combination of estrogen and progesterone were administered and animals were sacrificed after 12h to observe the effect of both steroid hormone alone as well as in combination. All tissues were harvested for further analysis. Only olive oil treated ovariectomized animals were considered as control.

### 2.7 Statistical analyses

Experimental data were collected from a minimum of five independent animals. Statistical analyses were done using Student’s *t-test* (for single comparison). For multiple comparison, data were analyzed by one-way ANOVA (analysis of variance) followed by Dunnett post hoc test [for comparison of peri-implantation period (from D2 to D5) vs. D1]. Data were considered statistically significant at p ≤ 0.05, and is indicated by an asterisk in the figures. All data was analyzed and plotted using GraphPad Prism 5.0 software and data were presented as mean ± SEM.

## 3. Results

### 3.1 Effect of anti-LIF antibody on embryo implantation

LIF is regarded as a most important mediator of embryo implantation in mammals. To know the role of this cytokine in embryo implantation of golden hamster, we used anti-LIF antibody to check the expression of LIF during window of implantation. We found significantly a smaller number of implanted embryos in anti-LIF antibody treated uterine horn as compared to control uterine horn (Figure 1) (P ≤ 0.05, Student’s *t-test*).

**Figure 1:**
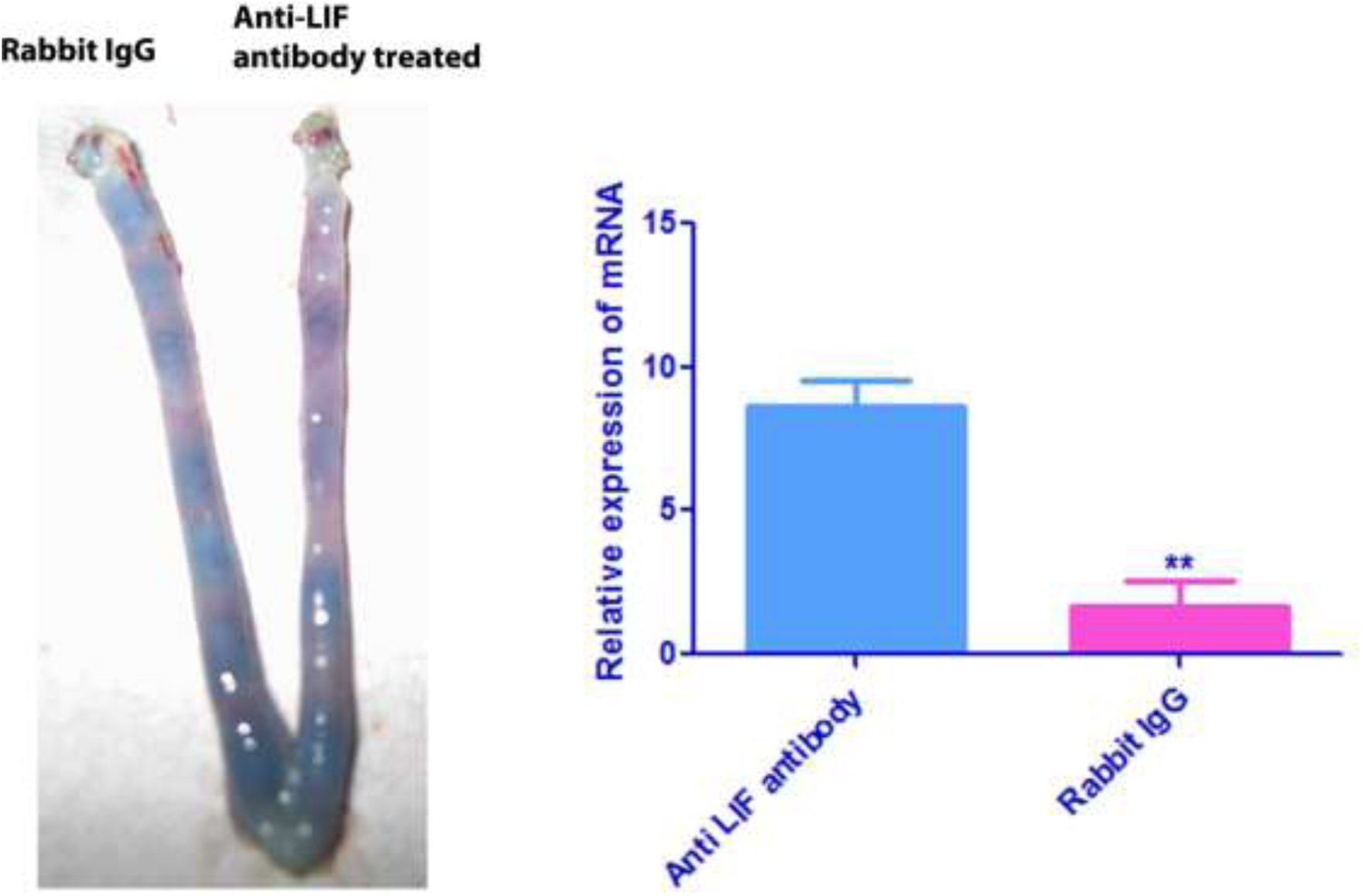
Effect of anti-LIF antibody on embryo implantation. Photograph of uterine horn of day 5 pregnant golden hamster showing the effect of intrauterine treatment of anti-LIF antibody. Treated animals were sacrificed on day 5 of pregnancy at 1000 hrs and implantation sites were observed using intravenous injection of pontamine sky blue dye. Significantly a smaller number of implantation site was observed in treated horn (P ≤ 0.05, Student’s *t-test*). Vehicle treated control horn, however, showed normal implantation sites.

### 3.2 Cell specific localization of LIF in uterus during early pregnancy and pseudopregnancy

In order to know the cell specific localization of LIF protein in the uterus of golden hamster during early pregnancy, we performed immunohisotochemistry in the uterus of different day of pregnancy. We observed the localization of LIF protein in pregnant uterus from day 1(D1) to day 7(D7) of pregnancy in cell specific pattern (Figure 2). LIF is mainly present in gladular epithelium and perimetrium during first and second day of pregnancy. Small amount of LIF was also observed in luminal epithelium and stromal cell in uterus of D1 and D2 pregnant hamster. Suddenly, a sweeping increase in immunopositive cell was observed in luminal epithelium, glandular epithelum and stromal cell of D3 and D4 of pregnancy which are normally considered as a time frame for window of implantation. LIF was observed strictly localized in luminal epithelium around the blastocyst at D5 of pregnancy. However, some amount of immunopositive cells is also found to be present in stromal cell in antimesometrial end of uterus in D5. In this day, mesometrum and perimetrium region of uterus contain almost negligible amount of immunopositive cell for LIF. We also observed the presence of LIF protein in stromal cells of endometrium along with luminal epithelium during D6 and D7 of pregnancy in which stromal cells modified in to decidual cell in a process of decidualization under the influence of steroid hormones.

**Figure 2:**
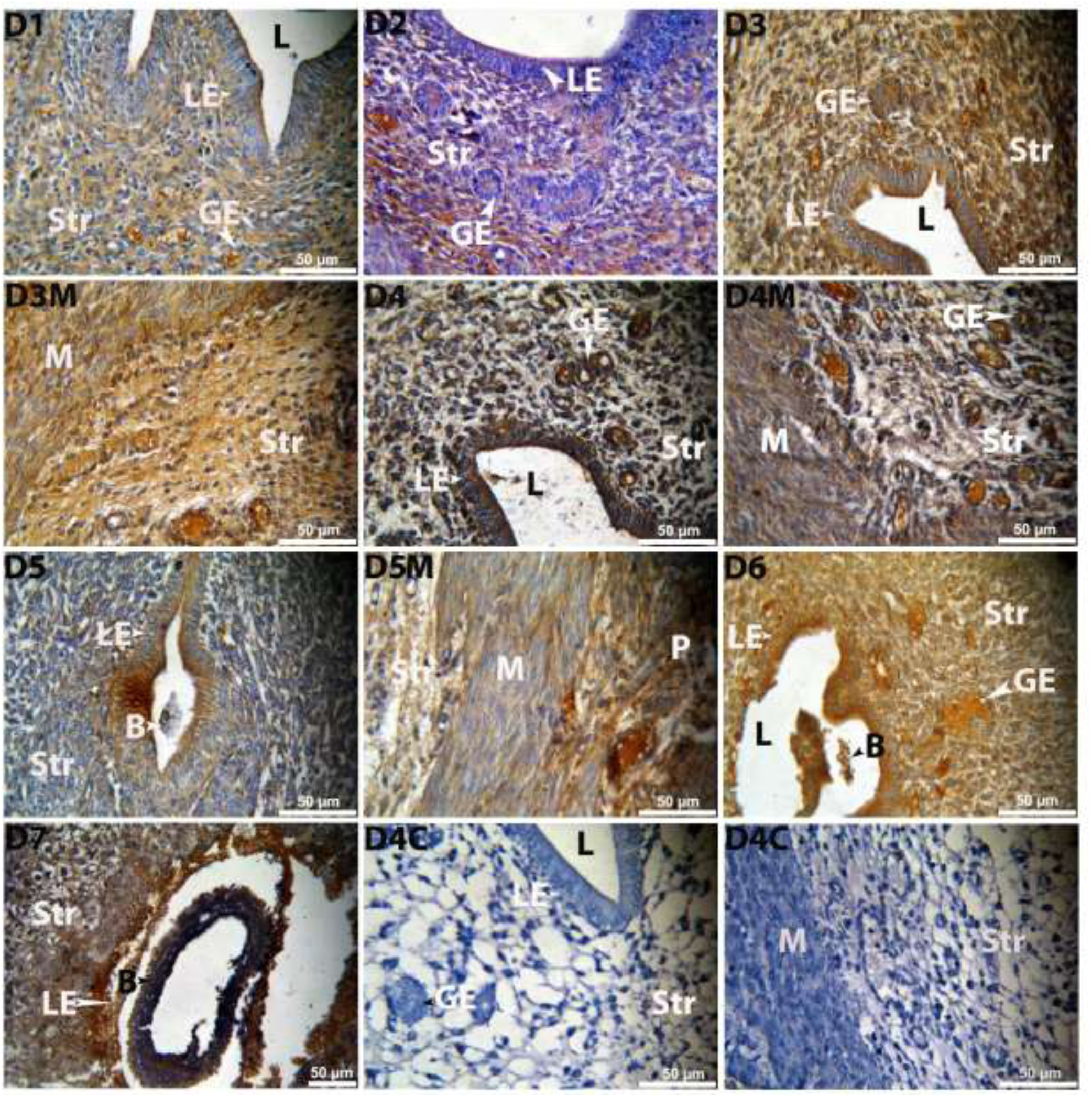
Immunolocalization of LIF in uterus during peri-implantation period. Immunohistochemical localization of LIF in stroma (Str), luminal epithelium (LE) and glandular epithelium (GE) of pregnant hamster uterus during early pregnancy (Day 1-7). Uterine horns were collected on day 1 to day 4 of pregnancy. D5 onwards, Implantation sites were separated. Rabbit polyclonal antibody is used for the detection of LIF antigen. However, negative control (D4C) was incubated with rabbit IgG. Positive staining showed brown colour. Photographs were taken under bright field condition. Magnification-X400, scale bar-50 µm and n = 5 in all the groups. Bl represents blastocyst and D represent day of pregnancy.

Successful pregnancy is a outcome of two-way communication between blastocyst and receptive uterus. All molecules required at physiological level to prime the uterus for blastocyst attachment is secreted by endometrium or /and blastocyst during early pregnancy. Many molecules are supposed to be synthesized by receptive endometrium under the influence of competent blastocyst. Therefore, it became necessary to know that whether the expression of LIF in the uterus of golden hamster during different day of pregnancy is depends on the presence of blastocyst. To answering this question, we prepared a pseudopregnant model of golden hamster by mating between normal cyclic female and vasectomized reproductively stud male. The hormonal milieu and microenvironment in uterus are exactly similar to that in pregnancy except the presence of true blastocyst. We found almost similar pattern of expression of LIF protein in the uterus of pseudopregnant hamster as in pregnancy (Figure 4A). Significant amount of immunopositive cells for LIF was observed in luminal epithelium and stromal cell of D3 and D4 of pseudopregnancy. Stromal cells of D5 pseudopregnant hamster also contain LIF positive cells in luminal epithelia with a smaller number of immunopositive cells in stroma as compared to the uterus of D3 and D4 pseudopregnant hamster. Onwards D5, the number of LIF immunopositive cell decreases with least visible in D6 and D7 of pseudopregnancy.

### 3.3 Gene expression analysis of LIF in uterus during early pregnancy and pseudopregnancy

To know the transparent role of LIF during embryo implantation and decidualization, we investigated their expression in uterus during normal pregnancy and pseudopregnancy at mRNA level by quantitative and semiquantitative reverse transcriptase PCR using hamster specific primers (Table-1 and 2). We observed that relative expression of their mRNA is significantly high in all the day during early pregnancy but their fold change varies in different day as compared to D1 (Figure 3A). Their expression is found to be upregulated around window of implantation with highest expression in D3 as compared to D1. mRNA of LIF was also found to be expressed in D6 to D8 of normal pregnancy. The relative expression of mRNA in the uterus of D7 of normal pregnancy was not found to be significantly differ as compared to D6 while it gets significantly decreased in D8 of pregnancy (Figure 3B).

**Figure 3:**
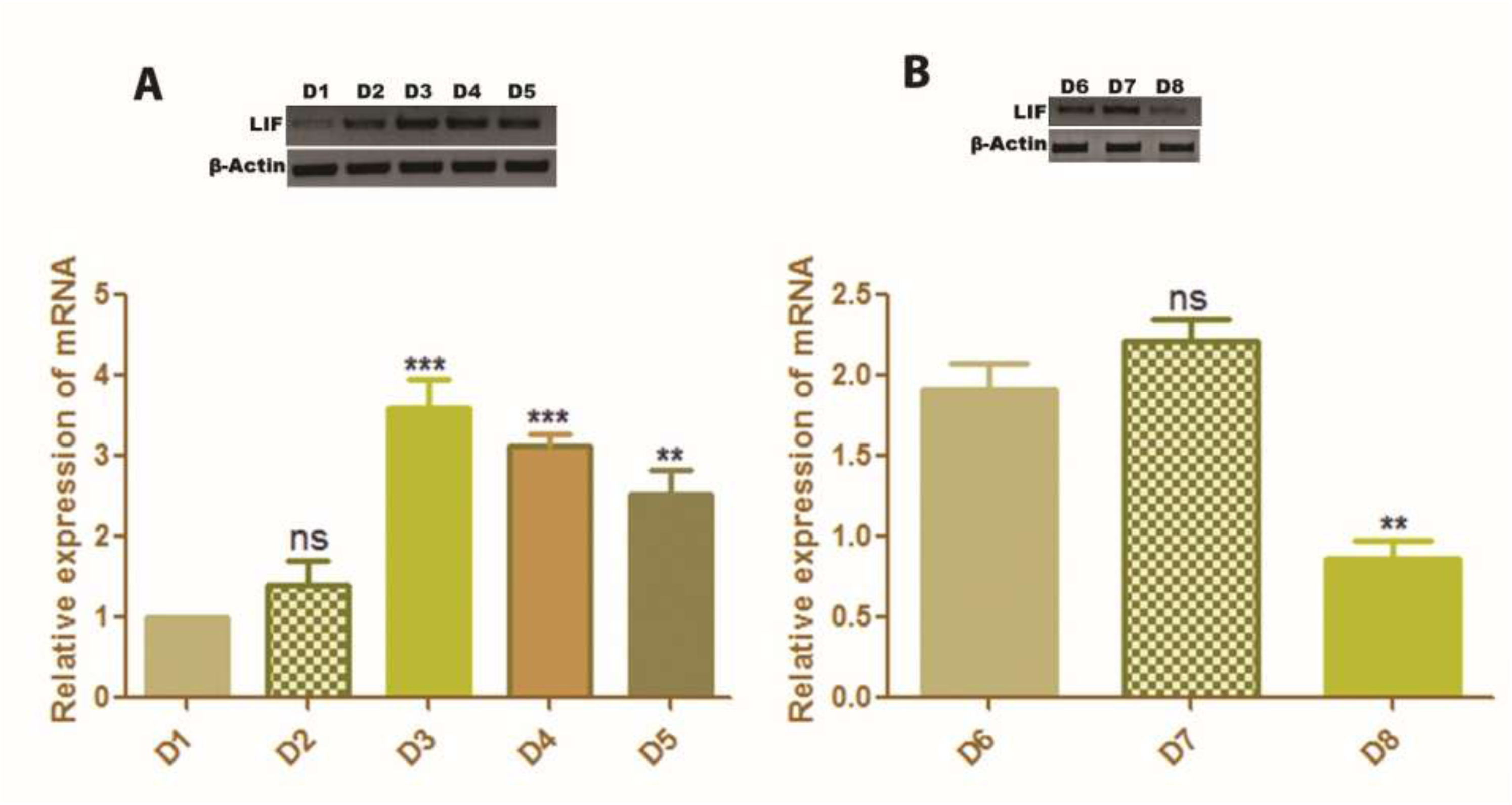
Gene expression analysis of LIF mRNA. Relative level of expression of mRNA (gel image for semiquantitative PCR and Graph for Quantitative PCR) of LIF mRNA in the uterus of golden hamster during early pregnancy were measured by real time and reverse transcription PCR. Total RNA was extracted from uterine horn of pregnant hamster using Trizol reagent, reverse transcribed using c-DNA synthesis kit and performed real time quantification using SYBR Green Supermix. **A.** LIF mRNA during pre and peri-implantation period (D1 to D5) (B) LIF mRNA during post implantation period (D6 to D8). Fold changes are expressed in relative to day 1 of pregnancy in pre and peri-implantation while that in post implantation period were expressed relative to day 6 of pregnancy. Data represent the mean ± SEM of three independent biological experiments. P value ≤ 0.05 is considered significant (* for P ≤ 0.05, ** for P ≤ 0.01 and *** for P ≤ 0.001).

**Figure 4:**
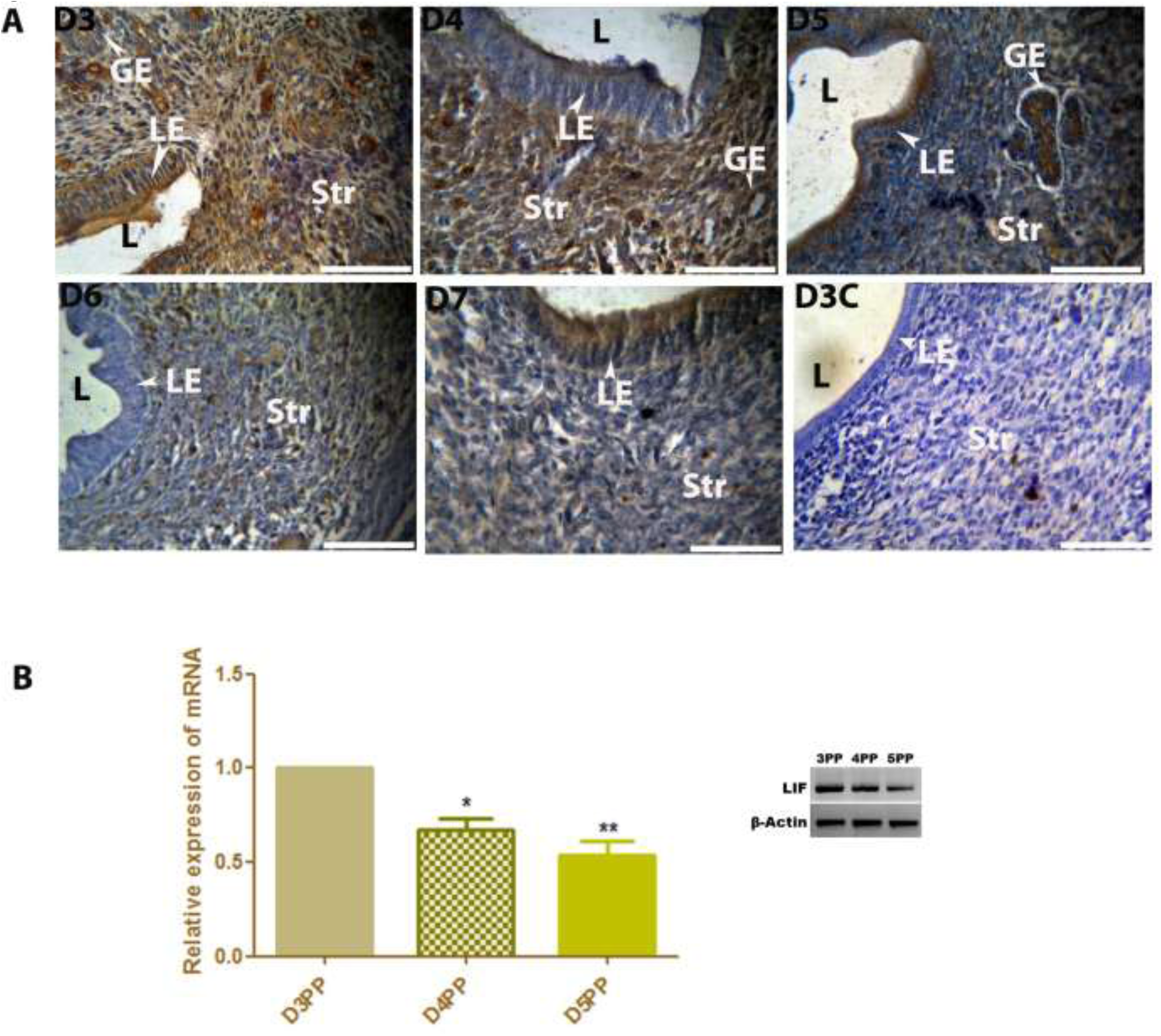
**A.** Immunohistochemical localization of LIF protein in stroma (Str), luminal epithelium (LE) and glandular epithelium (GE) of pseudopregnancy (D3 to D7 pseudopregnancy). **B.** Gene expression analysis of LIF mRNA (gel image for semiquantitative PCR and Graph for Quantitative PCR) in D3 to D5 of pseudopregnancy. PP in gel image represents pseudopregnancy of specified day. Fold changes are expressed in relative to day 3 of pseudopregnancy. Data represent the mean ± SEM of three independent biological experiments. P value ≤ 0.05 is considered significant (* for P ≤ 0.05, ** for P ≤ 0.01 and *** for P ≤ 0.001).

Uterus of pseudopregnant hamster also expressed LIF mRNA during early pseudopregnancy. However, relative mRNA expression in D3 pseudopregnancy is found to be upregulated as compared to other day of pseudopregnancy (Figure 4B).

### 3.4 Steroid hormones regulation LIF in uterus of ovariectomized female hamster

LIF is known to be an estrogen responsive cytokine in the uterus of mice during early pregnancy. Although, it is a well-established fact that golden hamster does not rely on ovarian estrogen for embryo implantation still we have checked their regulation by steroid hormone in the uterus of ovariectomized hamster. We observed that the expression of mRNA is upregulated in estrogen and progesterone alone as well as in combination of both hormones treated ovariectomized uterus as compared to vehicle (Oil) treated uterus but the fold change in expression in response to estrogen treatment is slightly high as compared to other (Figure 5). However, we did not observe any significant difference in fold change amongst all hormones treated group.

**Figure 5:**
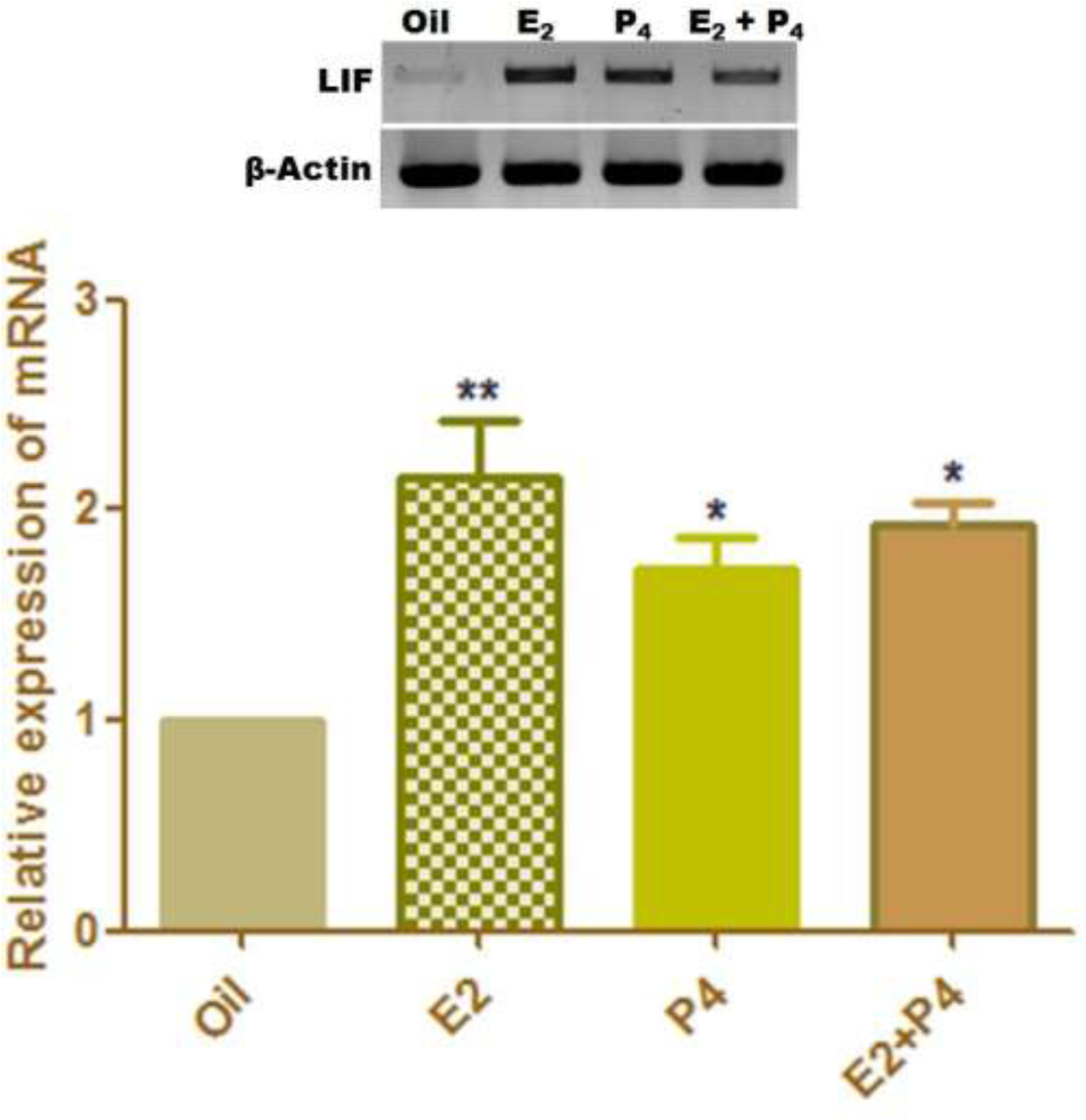
Hormonal regulation of LIF mRNA in the uteri of ovariectomized hamster (gel image for semiquantitative PCR and Graph for Quantitative PCR). E_2 -_estrogen, P_4_ - Progesterone and oil is representation of olive oil which was used as solvent for hormone. Fold changes are expressed in relative to oil treated (Vehicle) ovariectomized hamster. Data represent the mean ± SEM of three independent biological experiments. P value ≤ 0.05 is considered significant (* for P ≤ 0.05, ** for P ≤ 0.01 and *** for P ≤ 0.001).

### 3.5 Letrozole treated D5 uterus showed least expression LIF

Several implantation specific molecules are observed to express in uterus of mice and rat and found to be regulated by estrogen. These molecules are similarly found to express in uterus of golden hamster during window of implantation despite the absence of ovarian estrogen. It has been established that pregnant mice uterus is a novel site for the synthesis of estrogen and this uterine estrogen is sufficient to induce the signaling pathway associated with the process of decidualization and angiogenesis during early pregnancy (Das et al., 2009). Therefore, considering this hypothesis for hamster, we checked the expression of LIF in letrozole treated D5 (LD5) uterine horn. Letrozole is a inhibitor of an cytochrome P_450_ aromatase, a rate limiting enzyme for the biosynthesis of estrogen. Interestingly, we observed significantly less (t-test, p≤ 0.001) expression of mRNA of LIF in LD5 uterine horn as compared to (ND5) implantation site (Figure 6).

**Figure 6:**
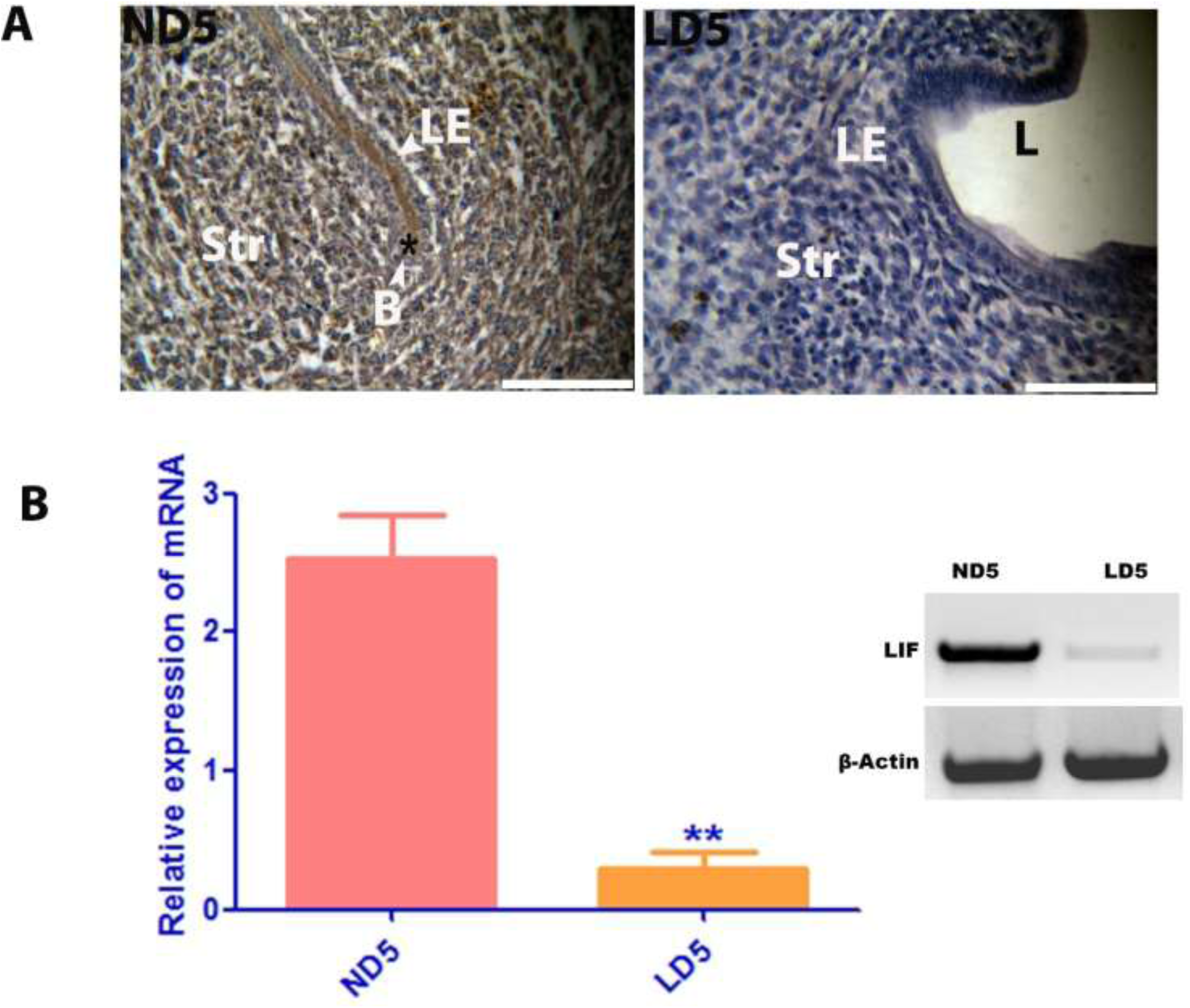
Relative expression of LIF mRNA (gel image for semiquantitative PCR and Graph for Quantitative PCR) in letrozole treated day 5 (LD5) uteri as compared to normal day 5 (ND5) pregnant uteri. Data represent the mean ± SEM of three independent biological experiments. P value ≤ 0.05 is considered significant (Student’s *t-test*).

### 3.6 Anti-LIF antibody treated uterine horn showed less expression of COX-1, COX-2, mPtges and mPtgis

prostaglandins are responsible for increased vascular permeability at the implantation site which is a prerequisite for successful embryo implantation in mice (Dey et al., 2004). Prostaglandins are synthesized by the activity of COX-1 and COX-2. Expression of these enzymes were also observed in the uterus of golden hamster during early pregnancy (Wang et al., 2004). But their upstream regulator during window of implantation in golden hamster is largely unknown. We checked the expression of these enzymes in the anti-LIF antibody treated uterine horn to know the possibility of any synergism between LIF and these enzymes. We observed that the expression of mRNA of COX-1, COX-2, mPtges and mPtgis were significantly reduced (P≤0.001, Student’s *t-test*) as compared to normal day 5 implantation site (Figure 7).

**Figure 7:**
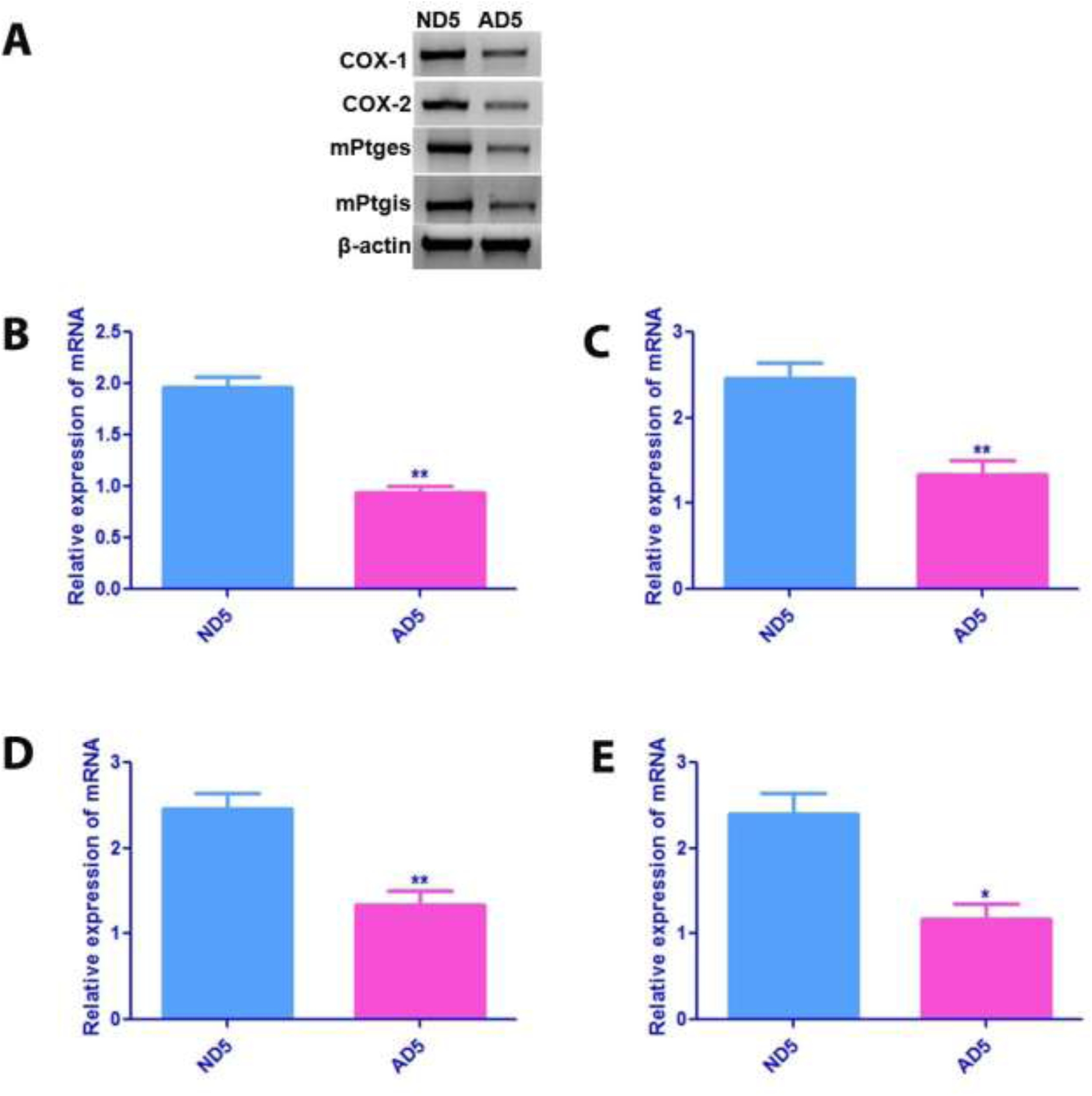
Relative expression of COX-1, COX-2, mPtges and mPtgis mRNA (gel image for semiquantitative PCR and Graph for Quantitative PCR) in anti-LIF treated day 5 uterine horn. Data represent the mean ± SEM of three independent biological experiments. P value ≤ 0.05 is considered significant (Student’s *t-test*).

## 4. Discussion

LIF is important proinflammatory cytokines which is recruited by endometrium of pregnant female and expressed in spatiotemporal manner in different tissue compartment of uterus during early pregnancy. Maximum expression of uterine LIF was found during embryo implantation which strengthens the critical significance of LIF in this process (Bhatt et al., 1991). Furthermore, it’s crucial role in implantation was demonstrated by the targeted deletion of LIF and its receptor in mice which results in abnormal embryo implantation and decidualization (Stewart et al., 1992). However, there are very few reports exist which explain the expression and regulation of LIF in golden hamster. The detailed study of LIF signaling in golden hamster enables us to understand the precise signaling mechanism of endometrial receptivity in mammals where implantation does not require ovarian estrogen akin to human. Our study is a tiny attempt in this context to know the regulation of LIF by nonsystemic estrogen during early pregnancy.

We observed significantly a smaller number of implantation site in anti-LIF antibody treated uterine horn as compared to control horn. This implies the importance of LIF in the uterus of golden hamster during window of implantation. Our results showed similarity with previous finding of Ding et al., 2008; in which they used slightly high dose of LIF antibody to determine any probable role of LIF in early pregnancy. Further, we checked the expression of LIF protein in the uterus of golden hamster during early pregnancy to know their involvement in the process of embryo implantation and decidualization separately. We found that LIF is largely localized in the luminal epithelia during early pregnancy and their expression is upregulated in D3 of pregnancy. Several reports are available which considered that the presence of LIF in luminal and glandular epithelia is essential for the successful event of implantation in many mammals (Shen et al., 1992; Cullinan et al., 1996, Bhatt et al., 1991). Our results is consistent with the previous finding in mice where LIF and their receptors are found to be expressed in luminal epithelia during window of implantation (Bhatt et al., 1991). Luminal epithelia of rat endometrium were also supposed to harbor LIF during embryo implantation. LIF present in luminal epithelium of uterus is assumed to be responsible for activating at least 50 transcription factors within a very short time of induction of LIF which modulate several physiological pathways that involves in successful event of embryo implantation and decidualization (Rosario et al., 2016; Rosario et al., 2014). This reflect the stunning ability of a single cytokines to induce a dynamic and complex network which is responsible for transforming hostile luminal epithelia in to hospitable which support the attachment of blastocyst. Any mutation in LIF or LIF receptor related gene in luminal epithelia results in inactivation of JAK-STAT pathway (Lee et al., 2013; Ernst et al., 2001; Sun et al., 2013). Furthermore, the inhibition of homeobox gene MSX1 expression in luminal epithelia is a prerequisite for the endometrial receptivity which is mediated by the presence of LIF in luminal epithelia. Persistent expression of homeobox MSX1 gene was supposed to responsible for the inactivation of JAK-STAT pathway in luminal epithelia (Daikoku et al., 2011) which ultimately leads to implantation failure as like in mice treated with JAK-STAT inhibitor before implantation (White et al., 2007). This phenomenon was also observed in monkey that became infertile due to abnormality in embryo implantation when treated with monoclonal antibodies of LIF (Ghosh et al., 2005). Although, blastocyst without functional LIF and/or their receptors showed normal implantation process when transferred in to pseudopregnant uterus. this reflects the importance of expression of LIF in luminal epithelia of pregnant uterus. Together these findings, suggest that luminal epithelia is the primary site for the action of LIF for successful embryo implantation. Therefore, we anticipate that upregulated expression of LIF in luminal epithelium just prior to implantation (D3 of pregnancy) might be associated with the activation of JAK-STAT pathway to make endometrium receptive for proper embryo implantation in golden hamster.

Glandular epithelium of uterus was supposed to play an important role in establishment of pregnancy in rodents (Cooke et al., 2012; Filant and Spencer, 2013b, Gray et al., 2001c). Cross section of uterine wall contains about 10-20 uterine glands surrounded by epithelial cells and predominantly present in the antimesometrial end of uterus where generally embryo get attached. Endometrial glands are found to be present in all mammalian uteri that are involved in synthesis and/or transport and secretion of essential substance for the survival and development of embryo (Filant et al., 2014). Our results demonstrate the presence of LIF in glandular epithelium of golden hamster. This finding corroborates the previous observation of Colin Stewart in which they observed the presence of LIF in the glandular epithelium of mice during early pregnancy in response to nidatory estrogen (Bhatt et al., 1991). Occurrence of LIF in glandular epithelium was also observed in other mammalian species including human (Shen et al., 1992; Cullinan et al., 1996). They are found to express maximally just prior to embryo implantation. The expression of LIF at mRNA level in glandular epithelial suddenly increases within 1 hr of estrogen induction and found to persists for further 5-6 hrs. Their expression in glandular epithelium was supposed to be regulated by tumor suppressor transcription factor P^53^ in mice along with estrogen receptor (Cheng et al., 2001). LIF present in glandular epithelia is secreted in to uterine lumen where it is supposed to binds with their receptor complex present on the apex of luminal epithelia and provide addition support for successful embryo implantation by phosphorylating JAK-STAT pathway (Hu et al., 2007). The existence of LIF in the glandular epithelia during the window of implantation in hamster might be responsible for enhancing embryo implantation through aforesaid mechanism.

We also detected the expression of LIF in uterus of golden hamster during pseudopregnancy. This finding is in line with the burst expression of LIF in uterus of D4 pseudopregnant mice (Bhatt et al., 1991). This implies that the onset of expression of LIF in uterus is independent from blastocyst and is induced by maternal factors. In mice, it is supposed that the high level of circulating estrogen during D4 of pregnancy and pseudopregnancy is responsible for the upregulated expression of LIF (Bhatt et al., 1991). Since, circulating estrogen derived from ovary does not play any important role in implantation in golden hamster, therefore, it could be possible that rise in expression of LIF in uterus during pregnancy and pseudopregnancy is induced by *de novo* synthesized estrogen in uterus during early stage of pregnancy like in mice (Das et al., 2009).

Our study revealed the expression of LIF in stromal cells immediate after implantation. Similar results has been also observed in mice and human in which LIF was supposed to involve in the decidualization of stromal cells for the support of developing embryo. This progression is found to be badly hampered by administration of LIF antagonist Shuya et al., 2011). The procedure of decidualization in LIF null mouse is overcome by administrating recombinant LIF from outside. This finding additionally supports the role of LIF in decidualization. The regulation of LIF in the luminal epithelia is mediated by canonical and non-canonical Wnt pathway (Rosario et al., 2014). Canonical Wnt/β catenine is supposed to be involved in the priming of uterus during early pregnancy before and after embryo attachment (Rosario et al., 2014; Mohamed et al., 2005). Mice deficient in stromal β catenine are found to be sterile due to defect in implantation and decidualization (Zhang et al., 2012). Wnt7A is a type of ligand, responsible for the nuclear translocation of β catenine in luminal epithelia as well as in stromal cells. The induction of this pathway is finally mediated by LIF by activating Wnt7A. Observed expression of LIF in stromal cells in golden hamster during post implantation (D5, D6 and D7) is probably associated with the successful decidualization following above mechanism.

Ovarian estrogen and progesterone are supposed to regulate the uterine LIF and their receptors during early pregnancy. Estrogen surge just prior to embryo implantation in mice is responsible for upregulation of LIF during window of implantation (Sherwin et al., 2004). We also observed almost similar results for the regulation of LIF in the uterus of ovariectomized hamster primed by different hormonal recipe. We found that estrogen and progesterone both are able to enhance the expression of LIF mRNA but the extent of expression due to estrogen only is significantly high as compared to other. Our results are in agreement with previous finding in golden hamster (Ding et al., 2008). However, they have been reported that LIF receptor is regulated by progesterone. We know that ovarian estrogen does not involve in embryo implantation in hamster and human. Therefore, we cannot decipher the signaling mechanism of LIF during embryo implantation in hamster and human on the basis of fact obtained from ovariectomized hormonal primed model. In this context, our results showed almost negligible expression of LIF in letrozole treated uterus during pregnancy. It clearly suggests that LIF during early pregnancy might be regulated by locally synthesized estrogen in uterus itself during implantation window. These results showed similarity with the regulation of LIF in mice and rat except their source which is ovary in mice and rat but might be uterus in case of golden hamster. These finding will give a new insight in future for the management of infertility in women.

Synthesis of prostaglandin by the activity of cyclooxygenases were also observed to induce by LIF during early pregnancy (Horita et al., 2007). We also observed the downregulation of cyclooxygenases along with mPtges and mPtgis in anti-LIF antibody treated uterine horn in golden hamster. This indicate the LIF mediated expression of prostaglandin in the uterus of hamster during early pregnancy. Our observation is also supported by the previous finding in mice where the expression of COX-2 was found to be downregulated in LIF null mice (Song et al., 2000; Kimber et al., 2005). The activity of prostaglandins was also observed to induce by IL-1β in mice (Nilkaeo and Bhuvanath, 2006). It has been demonstrated that LIF is responsible for inducing stromal cell decidualization by activating other cytokines and prostaglandins (Salleh et al., 2014).

Synergistic expression of COX-1, COX-2, mPtges and mPtgis in hamster uterus during early pregnancy might be associated with the process of decidualization to support further growrh and development of growing blastocyst.

## 5. Conclusion

Our results propose spatiotemporal expression of LIF in uterus of golden hamster during early pregnancy and pseudopregnancy which might be essential for successful event of embryo implantation and decidualization. Although, estrogen and progesterone are able to increase the expression of LIF in uterus of ovariectomized hamster, but almost negligible expression of LIF in D5 pregnant letrozole treated uterine horn could suggest their regulation by locally synthesized estrogen in uterus itself as a paracrine manner. It needs more investigation using knockout model to confirm it. Interdependent expression of prostaglandin synthesizing enzyme and LIF might be associated with the activation of inflammatory signals which are essential for successful establishment of early pregnancy in golden hamster.

## Acknowledgements

We are thankful to Banaras Hindu University, Varanasi, India (F (A)1-18 (H)/2531), for providing financial support for this work and Rockefeller Foundation, USA (P30/19567), for providing laboratory research infrastructure. We are also thankful to University Grants Commission (UGC) BSR (Zool/2011-12/39), India, for providing junior research fellowship (JRF) and Senior Research Fellowship (SRF) to Randhir Kumar.

The authors declare that there is no conflict of interest.

